# Impact of stroke on diurnal heart rate variability in rats

**DOI:** 10.1101/071555

**Authors:** Sean R. Huff, Samreen Ahmed, He Meng, Tiecheng Liu, Jimo Borjigin, Michael M. Wang

**Author notes:** Corresponding author: Michael Wang, 2215 Fuller Road, Veterans Affairs Building 31, Room 209, Ann Arbor, MI 48105-0532, **Email:**.

## Abstract

In humans, ischemic stroke is associated with reduced heart rate variability (HRV), a predictor of neurogenic cardiac events. This study was conducted to determine whether experimental stroke preferentially affects HRV during specific times of the day or parts of the sleep-wake cycle. A continuous telemetry system was used to analyze electroencephalograms (EEG), electromyograms (EMG), and electrocardiograms (ECG) after experimental ischemic stroke in rats (n = 13). Animals were implanted with telemetry probes and, after a two week stabilization period, 72h of baseline continuous EEG, EMG, ECG data were collected. Animals were then assigned to two procedure groups: 1) experimental stroke via 90 minute intraluminal filament occlusion of the right middle cerebral artery, or 2) sham surgery. Continuous waveform recording and HRV analysis was then continued for 48h following each procedure. HRV analysis was conducted by both time-domain and frequency-domain methods. Heart rate exhibited circadian rhythms among all animals prior to and following both procedures. Neither procedure contributed to significant heart rate changes. We did not find sleep state or circadian-dependent alterations in HRV in baseline recording. However, after stroke, animals exhibited elevated high frequency (HF) HRV (>10% increase, p<0.05) specifically during daytime waking periods. This contrasts with studies in humans that demonstrate reduced HRV after stroke, suggesting that age, medical comorbidities, or species differences modify the overall response to cortical infarction of autonomic control centers.

## INTRODUCTION

Heart rate variability (HRV) is a marker of autonomic health and is frequently affected by disease states. For example, stroke patients exhibit a multitude of HRV alterations compared to healthy controls. Numerous studies highlight the ability of diverse cerebral lesions to decrease total HRV (Barron et al. 1994). Patients who die or exhibit worse functional outcomes after stroke often exhibit reduced HRV acutely and sub-acutely after stroke, suggesting an association between depressed HRV and poor outcome from stroke (Colivicchi et al. 2005).

Analysis of specific HRV components has proven useful in particular disease states including stroke. For example, low frequency (LF) heart rate variability is thought to reflect baroreceptor sensitivity and high frequency (HF) heart rate variability is thought to most reflect the magnitude of parasympathetic outflow onto cardiac tissues during respiratory sinus arrhythmia (Moak et al. 2005). In turn, the observation that stroke patients exhibit diminished LF and HF magnitude and/or abolished circadian variation of these variables suggests that autonomic dysfunction may predispose to or develop as a consequence of stroke (Tokgozoglu et al. 1999; Colivicchi et al. 2005).

A significant proportion of stroke patients (19%) experience fatal or serious cardiac events following stroke (Rincon et al. 2008). A fraction of these cardiac events include sudden cardiac death (SCD), defined as death that occurs within one hour of the development of symptoms suggestive of cardiac insult or injury. It is thought that post-stroke dysfunction of the autonomic nervous system, the principle modulator of HRV, may contribute to these events. In accordance with this hypothesis, the aforementioned depression of HRV among stroke patients may coincide with excessive depolarization and/or excitotoxicity of myocardial tissue secondary to aberrant signaling of the autonomic nervous system, resulting in fatal ventricular tachycardia, ventricular fibrillation, and other cardiac abnormalities (Meglic et al. 2001; Sander et al. 2001; Taggart et al. 2011).

This study addresses two limitations of HRV studies in humans with stroke. First, it is difficult to conclude that stroke directly causes, rather than merely correlates to, heart rate variability changes in stroke patients (Colivicchi et al. 2005; Kuriyama et al. 2010). This is due to the inability to obtain baseline HRV data prior to stroke in humans. Second, the timing of HRV alterations after stroke has not been fully defined in humans. Determining whether stroke affects HRV during a specific time of the day or during specific phases of the sleep-wake cycle has not been possible in human studies. Confounding factors in clinical studies include variable patient characteristics, stroke etiology, and infarct location (Tokgozoglu et al. 1999; Meglic et al. 2001). Equally important, human stroke occurs at various time of the day, and patients are subjected to both hospital care that inherently disrupts normal sleep as well as lighting that interferes with normal circadian rhythms.

To overcome these two limitations, we investigated the effects of stroke on HRV in animals. The ability to measure baseline HRV and then to experimentally induce stroke allowed us to test if stroke causes alterations in HRV (Tokgozoglu et al. 1999; Colivicchi et al. 2005). Stroke was induced during the daytime phase of the circadian cycle in young animals of the same strain. Furthermore, multimodal telemetric and physiological analysis in free-running rats housed in strictly regulated lighting conditions permitted determination of whether stroke-induced changes in HRV varied by sleep-wake sate or circadian timing. To induce stroke, we use a model of right middle cerebral artery occlusion (MCAO) previously established in the adult rat. Occlusion of the right middle cerebral artery was of particular interest because the right insular cortex is thought to be responsible for regulating sympathetic tone, the branch of the autonomic nervous system most frequently implicated in arrhythmogenesis (Meyer et al. 2004; Taggart et al. 2011). Moreover, multimodal physiological analysis was used to determine whether changes in HRV varied by sleep-wake state or circadian conditions.

## METHODS

### Animals

All animals were housed at ambient temperature under 12:12 light:dark conditions. Animals were implanted with 4ET radiotelemetric physiological monitors (DSI) that allowed capture of digital electroencephalography, electromyography, and electrocardiography data at a sampling frequency of 100 Hz in freely moving animals. Animals were allowed to recover from implantation surgery and then physiological data were obtained for up to 7 days. One group of animals (n = 6) was subjected to ischemic stroke by transient MCAO for 2 hours. A second group of rats (n = 7) was subjected to sham surgery that consisted of inserting a suture into the carotid artery without advancement of the suture into the brain. Some sham animals (n=3) underwent a brief anesthesia procedure 14 days prior to sham procedure so that the effects of anesthesia on HRV could also be quantified. Radiotelemetric monitoring was halted during surgery (1-2h), but animals were again monitored for two consecutive 24h periods beginning at 1800h on the date of their respective procedures. Some of the animals described here were used in a previous study which reported the effects of stroke on sleep in Sprague Dawley rats (Ahmed et al. 2011).

MCAO was performed, as previously described, 14-28 days after implantation of physiological monitoring probes; all surgeries took place between 09:30 and 15:30, during the animal’s light cycle (Ahmed et al. 2011). Rats were allowed to awaken and, following confirmation of neurological deficits (circling to the left or gait to the left), were then reanesthetized for removal of the occluding filament 90 minutes following its placement. At the end of the recording period, animals were euthanized and brains removed for histological analysis of infarcts using tetrazolium chloride staining (Figure 2). Animals that did not survive the 48h period following stroke were excluded from the study.

### Sleep stage and heart rate variability analysis

Pre- and post-procedure sleep-wake states were scored using previously published definitions (Ahmed et al. 2011). A normalization procedure was also undertaken to account for interindividual differences in EEG (Baracchi et al. 2008).

Time-domain and frequency analysis of HRV was conducted using Dataquest ART software (DSI, Minnesota, USA). Baseline heart rate variability activity was determined by analyzing 72h of continuous ECG data immediately prior to the date of procedure. Heart rate variability analysis was also completed on the continuous ECG stream acquired during the 48h period following stroke, sham, and anesthesia procedures. Epoch duration of 10s was selected to match sleep stage analysis.

Time-domain analysis of heart rate variability was conducted by calculating the standard deviation of normal R-R (“N-N”) intervals during each 10s epoch. Automatic waveform detection and marking was conducted using Dataquest ART software. Visual inspection and mathematical analysis of waveform markings was completed to ensure that ectopic beats or unmarked QRS complexes did not affect calculations of heart rate or interbeat interval standard deviation (IBISTD). We removed epochs (8-20%) that contained ectopic beats or unmarked QRS complexes.

Frequency analysis of HRV was conducted on the same epochs by calculating the spectral density of R-R interval plots. The power bands analyzed included LF (0.06 – 0.6 Hz), HF (0.6 – 2.4 Hz), and LF/HF. LF is thought to reflect mostly baroreceptor sensitivity in the form of cardio-vagal gain, while HF is thought to reflect parasympathetic cardiac influences in the form of respiratory sinus arrhythmia (Meyer et al. 2004; Moak et al. 2005; Goldstein et al. 2011). Following 120 Hz interpolation of the R-R signal, a modified FFT with Hanning window was used to calculate the normalized spectral density of each frequency band.

LF activity in the rat is largely attributed to baroreceptor-mediated Meyer waves that operate at a frequency of ∼0.4 Hz, completing a full waveform only 4 times during a 10s epoch (Rajendra Acharya et al. 2006). Therefore, it was necessary to conduct parallel LF and HF analysis using extended epochs of 300s to establish the efficacy of using an epoch length of 10s. Ectopic beats were eliminated from data analysis using a previously described method (Cai et al. 2006). Utilizing extended epochs also allowed us to conduct spectral analysis of the very low frequency (VLF) band ( 0.0 – 0.06 Hz), a component of heart rate variability thought to be mediated by transient humoral influences and other pulsatile factors (Berntson et al. 1997).

### Statistical analysis

Daytime epochs were defined as all epochs between 06:00h and 18:00h, whereas all others were considered nighttime epochs. Epochs were further marked and categorized based on sleep-wake state (‘night-REM’, ‘day-REM’, *et cetera*). All 10s epochs of a single animal with the same combination of sleep-wake state and circadian condition were used to calculate mean values for heart rate variability. These values were calculated for three periods: 1) The 72h period prior to the day of stroke or sham procedure and, 2) the first and second consecutive 24 hour periods following 1800h on the procedures. Because 300s epochs could not be matched to one sleep-wake state, 300s epochs were only sorted on the basis of day/night conditions.

A normalization procedure to account for inter-animal differences was utilized, calculating the ratio of mean values during the two periods following procedure (numerator) to mean values before procedure (denominator). Unless otherwise indicated, Mann-Whitney U tests were used to compare relevant variables across different states, experimental groups, and after procedures.

To mitigate the effects of multiple comparisons, linear models with binary ‘dummy’ variables were utilized to simultaneously analyze the entirety of anesthesia, sham, and stroke data. Specifically, binary variables pertained to the presence or absence of a given experimental procedure, sleep-wake state, or circadian condition. In turn, each linear model attempted to quantify the effect of binary explanatory variables on a specific HRV component.

## RESULTS

### Pre-Procedure Analysis

Baseline measurements in rats prior to stroke or sham surgery revealed robust nocturnal heart rate increases (α < 0.05) that were conserved during all sleep-wake conditions (Hashimoto et al. 1999). Heart rate did not vary between sleep-wake states (α > 0.05) during the night or the day. No statistically significant differences in heart rate variability were observed between circadian conditions and sleep-wake states.

### Heart Rate following Experimental Procedures

The effects of stroke or sham procedure on heart rate during specific sleep-wake and circadian states were investigated by comparing heart rate values before and after each procedure; comparisons were made for the three sleep/wake states (REM, NREM, and wake) during two circadian phases (day or night) to yield 6 total comparisons per day. These comparisons were repeated for two days, yielding a total of 12 heart rate comparisons. Animals exposed to sham or stroke procedures demonstrated daytime-specific increases in heart rate following the experimental procedure. For sham animals, 5 of 12 sleep-wake/circadian-specific comparisons exhibited statistically significant heart rate increases after the procedure. Among animals subjected to stroke, 3 of 12 sleepwake/circadian comparisons exhibited statistically significant heart rate changes relative to baseline. Overall, we did not find systematic differences in heart rate resulting from stroke (compared to sham).

### HRV following Experimental Procedures

Stroke induced changes in HRV measures that were not seen after sham procedure. Tables 1 and Figures 1 show HRV values among animals subjected to stroke and sham procedures. After normalization to pre-procedure values, animals undergoing stroke exhibited statistically significant (α < 0.05) elevation in HF specifically during wake-daytime conditions (Table 1; Figure 1). No changes in HF were observed during nighttime conditions. Lastly, no VLF changes were observed in any experimental group (data not shown).

**Fig. 1.**
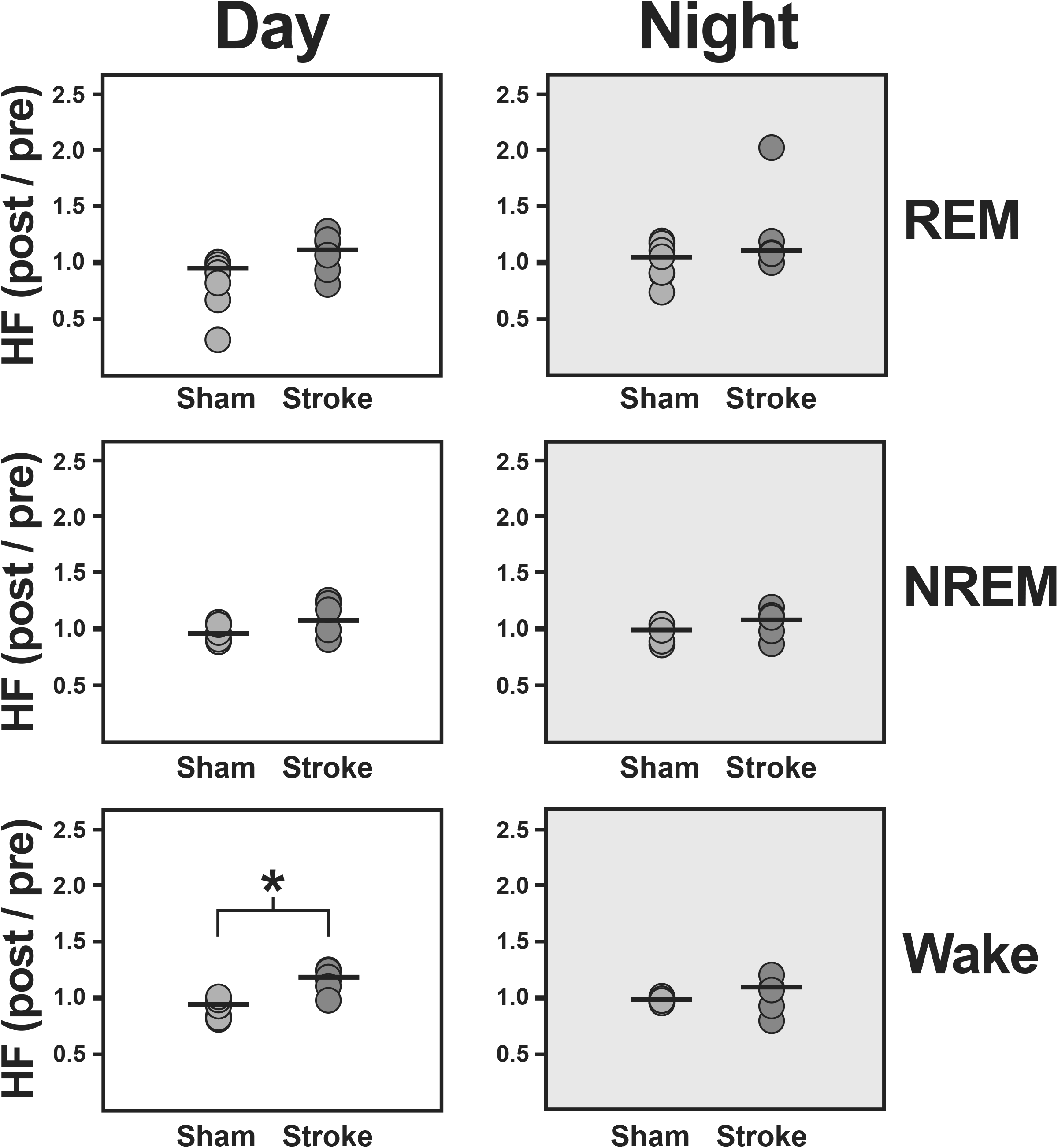
Scatterplot of HF changes following each procedure. The data was normalized by calculating the ratio of mean values during the second 24 hour period following each procedure (numerator) to mean values during the 72h period prior to the date of procedure (denominator). Circles indicate individual data points; black bars indicate group-wide averages. Asterisks indicate statistically significant between-group differences (p < 0.05)

**Tab. 1.** Normalized values for LF HRV (0.06 – 0.6 Hz), HF HRV (0.6 – 2.4 Hz), LF/HF ratio, and IBI-STD deviation of sham (n = 7) and stroke animals (n = 6) are shown for the, REM, and NREM stages of day and night conditions. Normalization consisted of calculating the ratio of mean values following each procedure (numerator) to mean values during the 72h period prior to the date of procedure (denominator). P-values were generated by a two-tailed t-test comparing normalized sham and stroke experimental group values during matched circadian and sleep-wake conditions. Emboldened and underlined values refer to normalized values among stroke animals that differ significantly (α = 0.05) from those of sham animals. Values within parenthesis indicate standard deviation

Post-Procedure HRV (Period 1)
|  |  | LF |  |  | HF |  |  | LF/HF |  |  | IBI-STD |  |  |
| --- | --- | --- | --- | --- | --- | --- | --- | --- | --- | --- | --- | --- | --- |
|  |  | Sham | Stroke | P-value | Sham | Stroke | P-value | Sham | Stroke | P-value | Sham | Stroke | P-value |
| REM | Day | **1.01**<br>(0.045) | **0.93**<br>(0.18) | 0.719 | **1.03**<br>(0.06) | **0.96**<br>(0.30) | 0.352 | **0.92**<br>(0.17) | **0.90**<br>(0.78) | 0.833 | **0.99**<br>(0.06) | **0.71**<br>(0.52) | 0.430 |
|  | Night | **0.96**<br>(0.048) | **1.00**<br>(0.22) | 0.617 | **1.04**<br>(0.07) | **0.98**<br>(0.18) | 0.352 | **0.98**<br>(0.16) | **0.90**<br>(0.44) | 0.226 | **0.99**<br>(0.12) | **1.37**<br>(1.00) | 0.285 |
| NREM | Day | **0.92**<br>(0.068) | **1.01**<br>(0.08) | 0.285 | **1.03**<br>(0.04) | **1.07**<br>(0.07) | 0.522 | **0.89**<br>(0.12) | **0.96**<br>(0.25) | 0.352 | **0.97**<br>(0.12) | **0.87**<br>(0.41) | 0.617 |
|  | Night | **0.98**<br>(0.050) | **0.98**<br>(0.13) | 0.944 | **1.00**<br>(0.03) | **1.07**<br>(0.12) | 0.174 | **0.96**<br>(0.18) | **0.92**<br>(0.35) | 0.944 | **0.99**<br>(0.09) | **1.20**<br>(0.48) | 0.352 |
| Wake | Day | **0.97**<br>(0.072) | **0.99**<br>(0.06) | 0.719 | <u>**1.05**</u><br>(0.03) | <u>**1.14**</u><br>(0.08) | <u>**0.027**</u> | **0.89**<br>(0.11) | **0.92**<br>(0.14) | 0.834 | **0.99**<br>(0.14) | **1.09**<br>(0.39) | 0.352 |
|  | Night | **0.95**<br>(0.060) | **0.99**<br>(0.09) | 0.522 | **1.00**<br>(0.03) | **1.15**<br>(0.16) | 0.352 | **0.94**<br>(0.12) | **0.81**<br>(0.31) | 0.429 | **0.98**<br>(0.09) | **1.49**<br>(0.51) | 0.352 |

Post-Procedure HRV (Period 2)
|  |  | LF |  |  | HF |  |  | LF/HF |  |  | IBI-STD |  |  |
| --- | --- | --- | --- | --- | --- | --- | --- | --- | --- | --- | --- | --- | --- |
|  |  | Sham | Stroke | P-value | Sham | Stroke | P-value | Sham | Stroke | P-value | Sham | Stroke | P-value |
| REM | Day | **1.00**<br>(0.08) | **0.99**<br>(0.16) | 0.834 | **0.94**<br>(0.11) | **1.17**<br>(0.19) | 0.073 | **1.09**<br>(0.44) | **1.09**<br>(0.27) | 0.352 | **1.01**<br>(0.19) | **1.60**<br>(0.71) | 0.332 |
|  | Night | **1.03**<br>(0.10) | **1.08**<br>(0.15) | 0.617 | **1.03**<br>(0.16) | **1.09**<br>(0.39) | 0.226 | **0.95**<br>(0.48) | **0.92**<br>(0.29) | 0.430 | **1.00**<br>(0.11) | **1.17**<br>(0.73) | 0.352 |
| NREM | Day | **1.03**<br>(0.18) | **1.04**<br>(0.80) | 0.617 | **0.96**<br>(0.08) | **1.10**<br>(0.14) | 0.174 | **1.24**<br>(0.39) | **1.29**<br>(0.22) | 0.617 | **1.03**<br>(0.09) | **1.31**<br>(0.59) | 0.719 |
|  | Night | **1.02**<br>(0.07) | **1.03**<br>(0.11) | 0.944 | **1.00**<br>(0.06) | **1.10**<br>(0.13) | 0.174 | **1.06**<br>(0.32) | **1.02**<br>(0.21) | 0.719 | **1.02**<br>(0.12) | **1.32**<br>(0.46) | 0.352 |
| Wake | Day | **1.03**<br>(0.13) | **1.00**<br>(0.08) | 0.617 | <u>**0.95**</u><br>(0.06) | <u>**1.17**</u><br>(0.11) | <u>**0.012**</u> | **1.17**<br>(0.26) | **0.99**<br>(0.20) | 0.101 | **1.01**<br>(0.12) | **1.26**<br>(0.47) | 0.134 |
|  | Night | **1.05**<br>(0.12) | **1.03**<br>(0.10) | 0.285 | **1.00**<br>(0.02) | **1.14**<br>(0.16) | 0.352 | **0.99**<br>(0.23) | **0.89**<br>(0.19) | 0.285 | **0.95**<br>(0.09) | **1.18**<br>(0.41) | 0.430 |

Linear models were used to verify statistically significant findings indicated by multiple two-tailed t-tests. Specifically, when sleep-wake state, circadian condition, and experimental procedure were considered binary variables, a linear model of HRV data revealed that stroke was the lone variable significantly contributing to or ‘driving’ HF and IBI-STD (coefficient p-values < 0.0001). Though the IBI-STD values of stroke animals did not differ from those of sham animals when using Mann-Whitney U test (Table 1), stroke contributed to increased magnitude of both HF and IBI-STD. A linear model of daytime epochs during the second period of post-procedure analysis indicated that stroke was the only statistically significant variable among a model of HF, exhibiting an overall R2 > 0.400. All other linear models (including those of the whole 48 hours of post-procedure analysis) exhibited R2 values between 0.100 and 0.400.

## DISCUSSION

This investigation explored the effects of right MCAO on heart rate variability in the rat during specific sleep-wake states and circadian times. Two main findings emerged: 1) Stroke elicited increased HF activity, and 2) Heart rate variability changes elicited by stroke occurred during waking periods only and were daytime-specific.

### Baseline heart rhythms, circadian rhythms, and sleep

At baseline, rats exhibited a nighttime heart rate elevation that is well-established within the literature (Hashimoto et al. 1999). The absence of influence of sleep-wake state on heart rate in animals at baseline contrasts with previous works that demonstrated depressed heart rate during sleep (del Bo et al. 1982). Our study design could account for some of the differences in our results; for example, we used outbred rats that may exhibit diverse physiological phenotypes. In addition, unlike earlier studies, our experiments used radiotelemetry devices in free-moving animals, eliminating the effects of an animal being tethered to its environment.

Post-surgical heart rate increases were observed among animals exposed to either stroke or sham procedures specifically during the day time. Such changes may reflect diffuse sympathetic activation among animals exposed to surgical stressors (Saad et al. 1989; Hachinski et al. 1992). The absence of heart rate differences during the nighttime may indicate the inability to detect pathologically elevated sympathetic activation during the period of physiological sympathetic dominance that occurs nocturnally (Hachinski et al. 1992).

### Daytime-specific increase in heart rate variability awake rats after stroke

Diminished total REM sleep in rats subjected to middle cerebral artery occlusion suggested that heart rate variability changes following stroke may vary with sleep-wake state (Ahmed et al. 2011). Indeed, rats subjected to stroke exhibited increased HF (a marker of cardiac parasympathetic influence) during the waking periods of daytime (Rajendra Acharya et al. 2006). The strict dependence of this HRV change on a sleep-wake state suggests that heart rate variability changes may be related to the same pathophysiological changes which drive previously demonstrated REM inhibition (Ahmed et al. 2011). But independent mechanisms that confine HRV changes to the day phase of the circadian cycle must exist, since REM sleep inhibition was observed during both the day and night (Ahmed et al. 2011).

We propose that a combination of sympathetic and parasympathetic upregulation with alteration of the baroreceptor reflex could explain our observations. In particular, dramatic sympathetic activation is known to take place following cerebral infarcts involving the insula (Meglic et al. 2001; Sander et al. 2001; Kuriyama et al. 2010). Anatomical studies confirmed infarction of this region in rats studied here (Figure 2). Previous investigations highlight the presence of cardio-protective vagal responses to elevated plasma norepinephrine (DeSilva et al. 1978; Saad et al. 1989). This parasympathetic response, mediated by the baroreflex, is primarily responsible for the elevation of HF and heart rate attenuation (Tulppo et al. 1998). The day-time specific timing of HF elevations may relate to the circadian baroreflex sensitivity observed in both rats and humans (Makino et al. 1997). Conversely, nocturnal sympathetic dominance with concurrent blunting of the baroreflex may limit the magnitude of compensatory parasympathetic outflow observed during nighttime epochs in the post-stroke period (Hashimoto et al. 1999; Meyer et al. 2004; Moak et al. 2005; Goldstein et al. 2011).

**Fig. 2.**
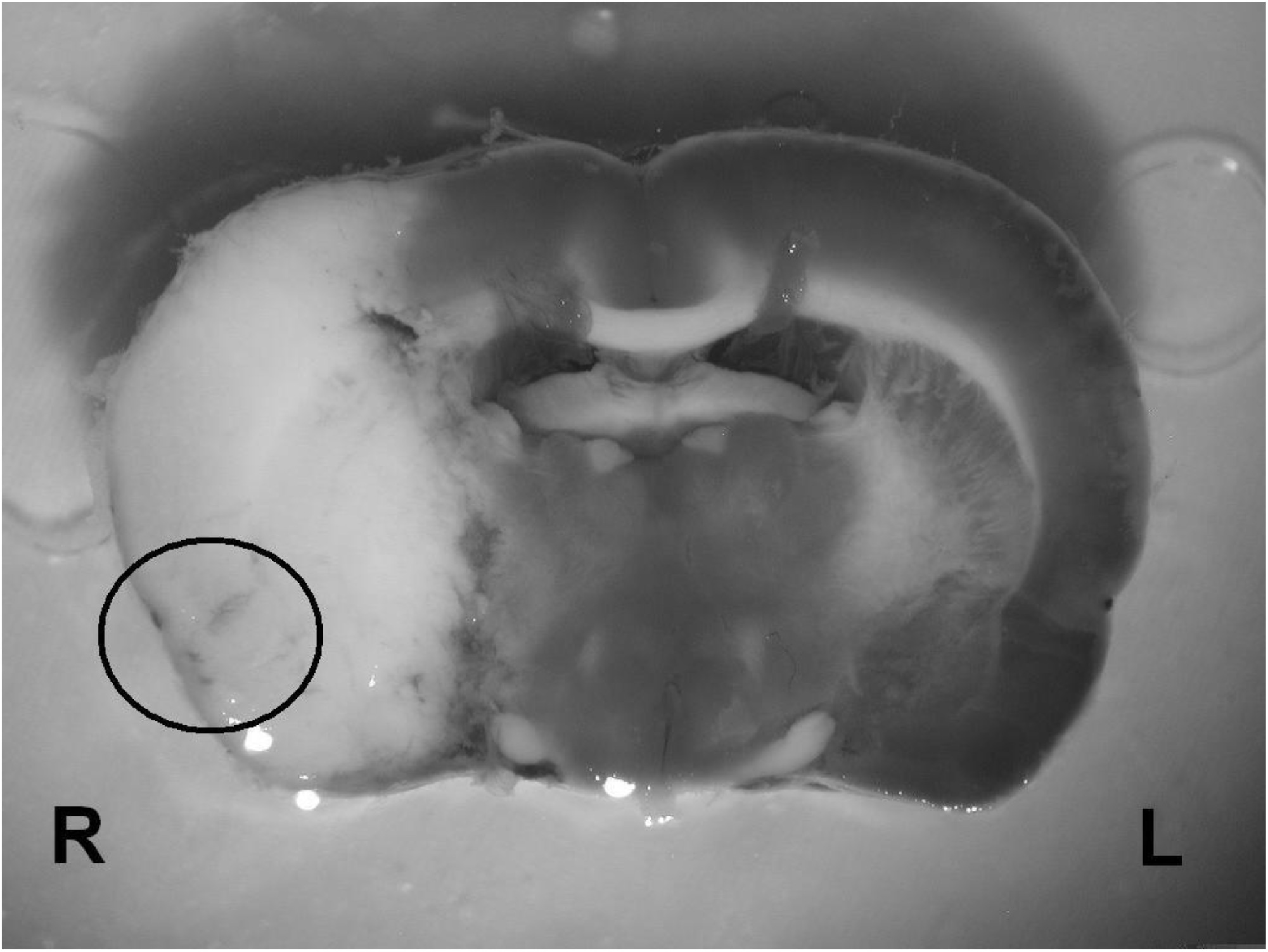
Rat brain after experimental stroke. The pale area represents infarcted tissue after tetrazolium chloride (TTC) staining; the encircled area within this larger pale area corresponds to insular cortex. ‘R’ indicates right-sided structures

Interestingly, HF has not been observed to increase in human stroke patients who are expected to have elevated plasma norepinephrine levels due to insular damage (Meglic et al. 2001; Sander et al. 2001; Kuriyama et al. 2010). The lack of HF elevation in the human during the post-stroke period may reflect baroreceptor insensitivity, manifesting as a blunted compensatory parasympathetic up-regulation. This interpretation supported by the findings that common stroke comorbidities (HTN, diabetes, aging) serve to inhibit baroreceptor signaling pathways and that baroreflex insensitivity also portends poor outcomes following stroke (Eveson et al. 2005; Heusser et al. 2005; Goldstein et al. 2011). Taken as a whole, these results suggest that the absence of parasympathetic up-regulation following stroke in human patients may portend poor prognosis in the form of unopposed sympathetic stimulation of heart tissue and a resulting increased likelihood of adverse cardiac event. Notably, our experiments did not use aged rats or rats with vasculopathy which clearly distinguish our preclinical model from published human studies. We also cannot exclude the possibility of cross-species differences in physiological changes after stroke, and it is also possible that factors such as timing of stroke onset, medications, and the disruptive effects of hospital care in could interfere with normal circadian cycles in humans, affecting measurement of diurnal changes in HRV.

In conclusion, we demonstrate that middle cerebral artery occlusion in the rat causes HF elevation. Changes in heart rate variability were specific to a single sleep-wake state and circadian condition: the wake state during daytime (Ahmed et al. 2011). Future investigations which manipulate the activity of the autonomic nervous system using pharmacological agents and also measure concentrations of norepinephrine will be useful to determine the specific mechanisms underlying our observations. Lastly, physically ablating or electrically stimulating the baroreceptor and testing the effects of ischemia in other brain regions could lead to strategies which enhance protective autonomic responses (such as daytime HF) that may prevent post-stroke cardiac excitotoxicity and arrhythmias (DeSilva et al. 1978; Taggart et al. 2011).

## ACKNOWLEDGEMENTS

We thank Mr. Yanming Li, Mr. Corey Powell, and Dr. Mark Opp for helpful discussions. Funding was provided by National Institutes of Health grant NS062816. We are also very grateful for the generous support of the Gilmore Fund for Sleep Research and Education.

## References

Ahmed S, Meng H, Liu T, Sutton BC, Opp MR, Borjigin J, Wang MM (2011) Ischemic stroke selectively inhibits REM sleep of rats. Exp Neurol 232:168-175 doi: 10.1016/j.expneurol.2011.08.020

Baracchi F, Zamboni G, Cerri M, et al. (2008) Cold exposure impairs dark-pulse capacity to induce REM sleep in the albino rat. J Sleep Res 17:166-179 doi: 10.1111/j.1365-2869.2008.00658.x

Barron SA, Rogovski Z, Hemli J (1994) Autonomic consequences of cerebral hemisphere infarction. Stroke 25:113-116

Berntson GG, Bigger JT, Jr., Eckberg DL, et al. (1997) Heart rate variability: origins, methods, and interpretive caveats. Psychophysiology 34:623-648

Cai Y, Qiu Y, Wei L, et al. (2006) Complex character analysis of heart rate variability following brain asphyxia. Med Eng Phys 28:297-303 doi: 10.1016/j.medengphy.2005.05.002

Colivicchi F, Bassi A, Santini M, Caltagirone C (2005) Prognostic implications of right-sided insular damage, cardiac autonomic derangement, and arrhythmias after acute ischemic stroke. Stroke 36:1710-1715 doi: 10.1161/01.STR.0000173400.19346.bd

del Bo A, Ledoux JE, Tucker LW, Harshkfield GA, Reis DJ (1982) Arterial pressure and heart rate changes during natural sleep in rat. Physiol Behav 28:425-429

DeSilva RA, Verrier RL, Lown B (1978) The effects of psychological stress and vagal stimulation with morphine on vulnerability to ventricular fibrillation (VF) in the conscious dog. Am Heart J 95:197-203

Eveson DJ, Robinson TG, Shah NS, Panerai RB, Paul SK, Potter JF (2005) Abnormalities in cardiac baroreceptor sensitivity in acute ischaemic stroke patients are related to aortic stiffness. Clin Sci (Lond) 108:441-447 doi: 10.1042/CS20040264

Goldstein DS, Bentho O, Park MY, Sharabi Y (2011) Low-frequency power of heart rate variability is not a measure of cardiac sympathetic tone but may be a measure of modulation of cardiac autonomic outflows by baroreflexes. Exp Physiol 96:1255-1261 doi: 10.1113/expphysiol.2010.056259

Hachinski VC, Wilson JX, Smith KE, Cechetto DF (1992) Effect of age on autonomic and cardiac responses in a rat stroke model. Arch Neurol 49:690-696

Hashimoto M, Kuwahara M, Tsubone H, Sugano S (1999) Diurnal variation of autonomic nervous activity in the rat: investigation by power spectral analysis of heart rate variability. J Electrocardiol 32:167-171

Heusser K, Tank J, Luft FC, Jordan J (2005) Baroreflex failure. Hypertension 45:834-839 doi: 10.1161/01.HYP.0000160355.93303.72

Kuriyama N, Mizuno T, Niwa F, Watanabe Y, Nakagawa M (2010) Autonomic nervous dysfunction during acute cerebral infarction. Neurol Res 32:821-827 doi: 10.1179/016164109X12464612122696

Makino M, Hayashi H, Takezawa H, Hirai M, Saito H, Ebihara S (1997) Circadian rhythms of cardiovascular functions are modulated by the baroreflex and the autonomic nervous system in the rat. Circulation 96:1667-1674

Meglic B, Kobal J, Osredkar J, Pogacnik T (2001) Autonomic nervous system function in patients with acute brainstem stroke. Cerebrovasc Dis 11:2-8 doi: 47605

Meyer S, Strittmatter M, Fischer C, Georg T, Schmitz B (2004) Lateralization in autonomic dysfunction in ischemic stroke involving the insular cortex. Neuroreport 15:357-361

Moak JP, Eldadah B, Holmes C, Pechnik S, Goldstein DS (2005) Partial cardiac sympathetic denervation after bilateral thoracic sympathectomy in humans. Heart Rhythm 2:602-609 doi: 10.1016/j.hrthm.2005.03.003

Rajendra Acharya U, Paul Joseph K, Kannathal N, Lim CM, Suri JS (2006) Heart rate variability: a review. Med Biol Eng Comput 44:1031-1051 doi: 10.1007/s11517-006-0119-0

Rincon F, Dhamoon M, Moon Y, et al. (2008) Stroke location and association with fatal cardiac outcomes: Northern Manhattan Study (NOMAS). Stroke 39:2425-2431 doi: 10.1161/STROKEAHA.107.506055

Saad MA, Huerta F, Trancard J, Elghozi JL (1989) Effects of middle cerebral artery occlusion on baroreceptor reflex control of heart rate in the rat. J Auton Nerv Syst 27:165-172

Sander D, Winbeck K, Klingelhofer J, Etgen T, Conrad B (2001) Prognostic relevance of pathological sympathetic activation after acute thromboembolic stroke. Neurology 57:833-838

Taggart P, Critchley H, Lambiase PD (2011) Heart-brain interactions in cardiac arrhythmia. Heart 97:698-708 doi: 10.1136/hrt.2010.209304

Tokgozoglu SL, Batur MK, Top uoglu MA, Saribas O, Kes S, Oto A (1999) Effects of stroke localization on cardiac autonomic balance and sudden death. Stroke 30:1307-1311

Tulppo MP, Makikallio TH, Seppanen T, Airaksinen JK, Huikuri HV (1998) Heart rate dynamics during accentuated sympathovagal interaction. Am J Physiol 274:H810-816

